# Protocol for intra-nerve AAV injection and dorsal root potential recording for optogenetic modulation of the peripheral sensory nerve activity

**DOI:** 10.64898/2025.12.10.693516

**Authors:** Akito Kosugi, Wupuer Sidikejiang, Shinji Kubota, Kazuhiko Seki

## Abstract

Optogenetic modulation of peripheral sensory nerve activity holds great potential for the treatment of sensory disorders. Here, we present a protocol for applying optogenetic techniques to peripheral sensory nerves using an adeno-associated virus (AAV) vector. We describe the procedure for gene transduction into dorsal root ganglion neurons via retrograde transport following intra-nerve AAV injection. We then outline a terminal, acute electrophysiological experiment to evaluate optogenetic effects at the level of the dorsal root.

For complete details on the use and execution of this protocol, please refer to Kosugi et al^1^.

**Graphical abstract:** 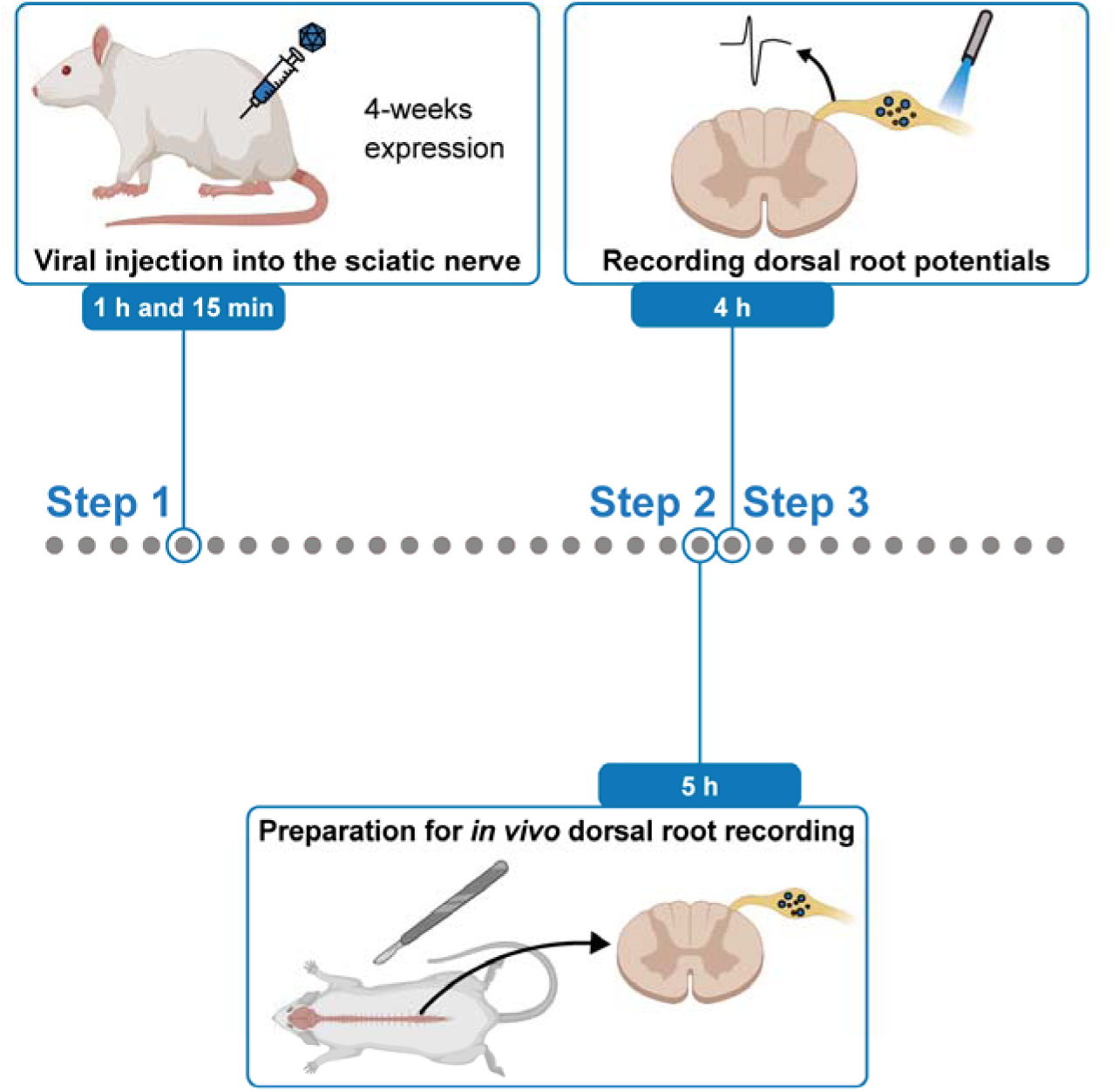

## Before you begin

The protocol below describes the specific steps for optogenetic modulation of peripheral sensory nerve activity. Due to the heterogeneity of peripheral sensory nerves, it is crucial to validate the functional types of sensory afferents modulated by optogenetics using electrophysiological methods. Compound action potentials recorded from the dorsal root of the spinal cord (i.e., dorsal root potentials) provide a reliable electrophysiological readout for assessing both the number and functional type of modulated afferent fibers. Here, we provide a comprehensive procedure for applying optogenetic manipulation to peripheral sensory nerves using an adeno-associated virus (AAV) vector, along with methods for precisely evaluating its effects. Specifically, we describe the injection of AAV into the sciatic nerve and the subsequent recording of dorsal root potentials at the lumbar level in male Wistar rats. This protocol can be readily adapted for other nerves and animal models, such as the common marmoset. For AAV injection procedures in marmosets, see Kudo et al^2^.

## Innovation

This protocol provides the first comprehensive methodology for reliably modulating peripheral sensory nerve activity using optogenetics. Although optogenetics has been widely established in the central nervous system^3,4^, its application to peripheral sensory nerves has been technically challenging due to the anatomical complexity and heterogeneity of dorsal root ganglion (DRG) neurons^5,6^.

Additionally, the lack of standardized electrophysiological readouts has limited the ability to quantitatively assess the effects of optogenetics. This protocol addresses these limitations.

First, the protocol employs an intra-nerve AAV injection strategy that enables robust retrograde transduction of DRG neurons while minimizing the procedural risks. Visualization of the viral solution with Fast Green dye provides immediate feedback on proper aspiration, unclogged ejection, and the absence of leakage, thereby ensuring consistent gene delivery across experiments.

Second, the protocol establishes dorsal root potential recording as a high-resolution, quantitative electrophysiological readout for evaluating optogenetic modulation. Because dorsal root potentials reflect the compound activity of primary afferent fibers, they enable the simultaneous assessment of both the number and functional type of modulated fibers, providing a standardized physiological measurement of the optogenetic effect.

Together, these improvements allow for the reliable modulation of peripheral sensory nerve activity using optogenetics. This provides a robust experimental framework for studying sensory processing and developing therapeutic strategies targeting peripheral nerve function.

## Institutional permissions

All surgery was performed according to the institutional guidelines for animal experiments and the National Institutes of Health Guide for the Care and Use of Laboratory Animals. All experiments were approved by the experimental animal committee of the National Institute of Neuroscience (approval number 2022020). Users must obtain all required institutional approvals before conducting any experiments described in this protocol.

### Preparation for viral injection Before the day of the experiment

Timing: > 1 week before

1. Prepare virus.

a. Purchase channelrhodopsin (ChR2) virus (AAV9-hSyn-ChR2(H134R)/EYFP) or halorhodopsin (eNpHR3.0) virus (AAV9-CMV-eNpHR3.0/EYFP).
b. Aliquot the virus and store at -80 °C.
2. Purchase animals.

a. Four-week-old male Wistar rats were purchased from vendors like Japan SLC, Inc.
3. Prepare the viral injection setup.

a. Order atipamezole, butorphanol, Fast Green, Fluorinert, medetomidine, midazolam, and lidocaine in accordance with institutional regulations.
b. Prepare surgery tools (see “Materials and equipment setup” section).
c. Prepare injection tools.

Timing: > 1 day before

1. Sterilize surgical tools using an autoclave.

### Day of the experiment

Timing: 1 hour

1. Prepare AAV solution.

a. Thaw the AAV solution aliquot on ice.
b. Dilute the AAV solution to the desired total volume (6 μL).

***Note*:** The AAV solution should be stored on ice.

a. c. Add 10% Fast Green to the AAV solution (0.6 μL) to enable visualization during injection. **CRITICAL:** Visualization of the AAV solution with Fast Green dye is important to confirm proper aspiration, unclogged injection, and leakage from the injection site.
b. 6. Prepare viral injection setup (Fig. 1A−C).
c. Connect a 31-gauge needle attached to polyethylene tubing (Fig. 1B) to a Hamilton syringe.

**Figure 1.**
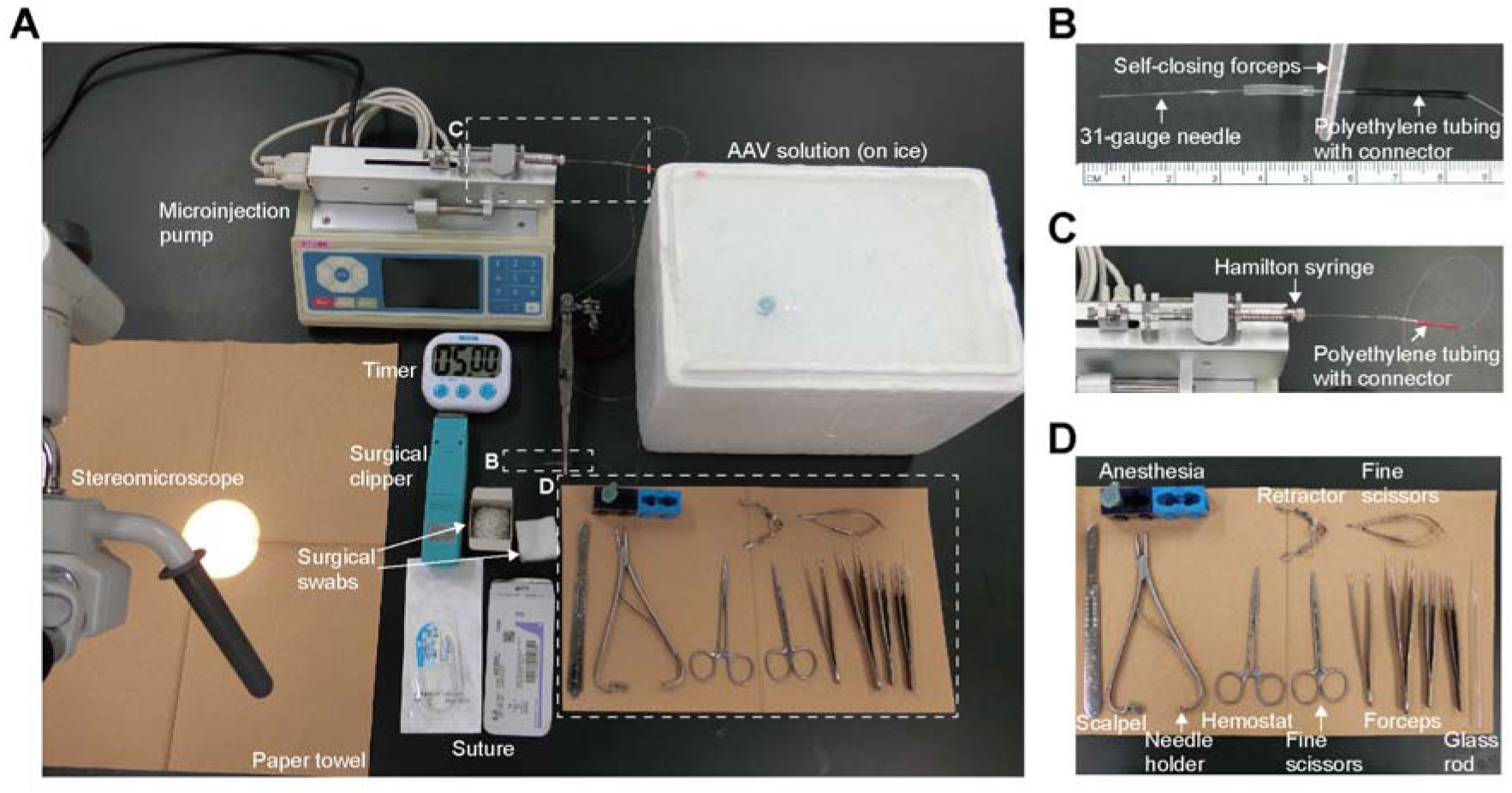
Viral injection preparation. (A) Surgical tools, anesthesia, and consumables for the viral injection. (B) Magnified view of the 31-gauge needle attached to polyethylene tubing, as indicated by the square in (A). (C) Magnified view of the Hamilton syringe mounted on a microinjection pump, as indicated by the square in (A). (D) Magnified view of the surgical tools, as indicated by the square in (A).

***Note*:** Connect the polyethylene tubing tightly; otherwise, the AAV solution cannot be aspirated.

a. b. Mount the Hamilton syringe on the microinjection pump (Fig. 1C).
b. Fill the syringe, tubing, and needle with an electrically insulating, stable fluorocarbon-based fluid (Fluorinert).
c. Withdraw the AAV solution into the needle and tubing, taking care to avoid introducing air bubbles.

***Note*:** During filling the syringe, avoid infusing too quickly, as it may increase internal pressure and cause the solution to continue leaking from the tip uncontrollably.

a. 7. Prepare the surgery setup, including surgical tools, anesthesia, and consumables including surgical swabs and sutures (Fig. 1A, D).

Preparation for the electrophysiological experiment Before the day of the experiment

Timing: > 1 week before

1. Prepare the electrophysiological recording setup.

a. Order isoflurane, mineral oil, paraformaldehyde, phosphate-buffered saline, rocuronium bromide, and thiopental sodium in accordance with institutional regulations.
b. Prepare surgery tools (see “Materials and equipment setup” section).
c. Prepare the electrodes (see “Materials and equipment setup” section).
d. Prepare a light source and a multimode optical fiber.

### Day of the experiment

Timing: 1 hour

1. Prepare the surgery setup, including surgical tools, anesthesia, and consumables including surgical swabs and sutures (Fig. 2A).
2. Prepare the respirator.

a. Check that there are no foreign materials inside the tracheal tube.
b. Confirm that the tip of the Y-shaped tube is appropriately shaped (Fig. 2B). If the tip is too long (greater than 10 mm) or the angle is too sharp, it may contact the tracheal wall and cause obstruction.
c. Verify that the ventilation rate (70 breaths per minute) and tidal volume (2.5 mL) are set appropriately.
3. Prepare the catheter for an intravenous line.

a. Check that there are no foreign materials inside the catheter.
b. Bevel the catheter tip at 45 degrees
c. Connect the catheter and the Cateran needle to a 10 mL syringe and fill it with saline.
4. 12. Prepare an electrophysiological recording.

a. In Clampex, select the “Acquire” tab and choose “New Protocol” from the list to create a new protocol, then save it as “volley recording”.

i. Set acquisition mode. Select the “Mode/Rate” tab and choose “Episodic stimulation” from the “Acquisition Mode” menu.
ii. Set sweep duration and sweeps per run. In the “Trial Hierarchy” menu, set “Trial delay (s)” to 0, “Sweeps/run” to 20, and “Sweep duration (s)” to 0.11.
iii. Set the sampling rate. In the “Sampling Rate per Signal” menu, set “Fast rate (Hz)” to 50000.
iv. Set the input channel. In the “Inputs” tab, check “Channel #0” checkbox, and select “IN 0” from the dropdown menu.
v. Set the trigger recording. In the “Trigger” tab, select “Immediate” from the dropdown menu under “Start trial with.”
vi. Select “Digitize START input” from the dropdown menu under “Trigger source.”
vii. Select “Start Sweep” from the dropdown menu under “Action.”
viii. Set “Timeout (ms)” to 1000.
b. Set the electrical stimulator parameters as follows: interval, 500 ms; set cycles, 20; delay, 10 ms; duration, 0.1 ms (electrical stimulation), 1 ms (optogenetic induction) or 500 ms (optogenetic suppression); train, 1.
c. Set the amplifier parameters as follows: sensitivity, 1 mV/V (×1000); low cut, 15 Hz; high cut, 10 kHz.
d. Set the light source. In “Doric Neuroscience Studio,” set the configuration to deliver light upon external trigger input, and adjust the intensity to 1000 mA. Set a 593-nm filter on the light source as well.

**Figure 2.**
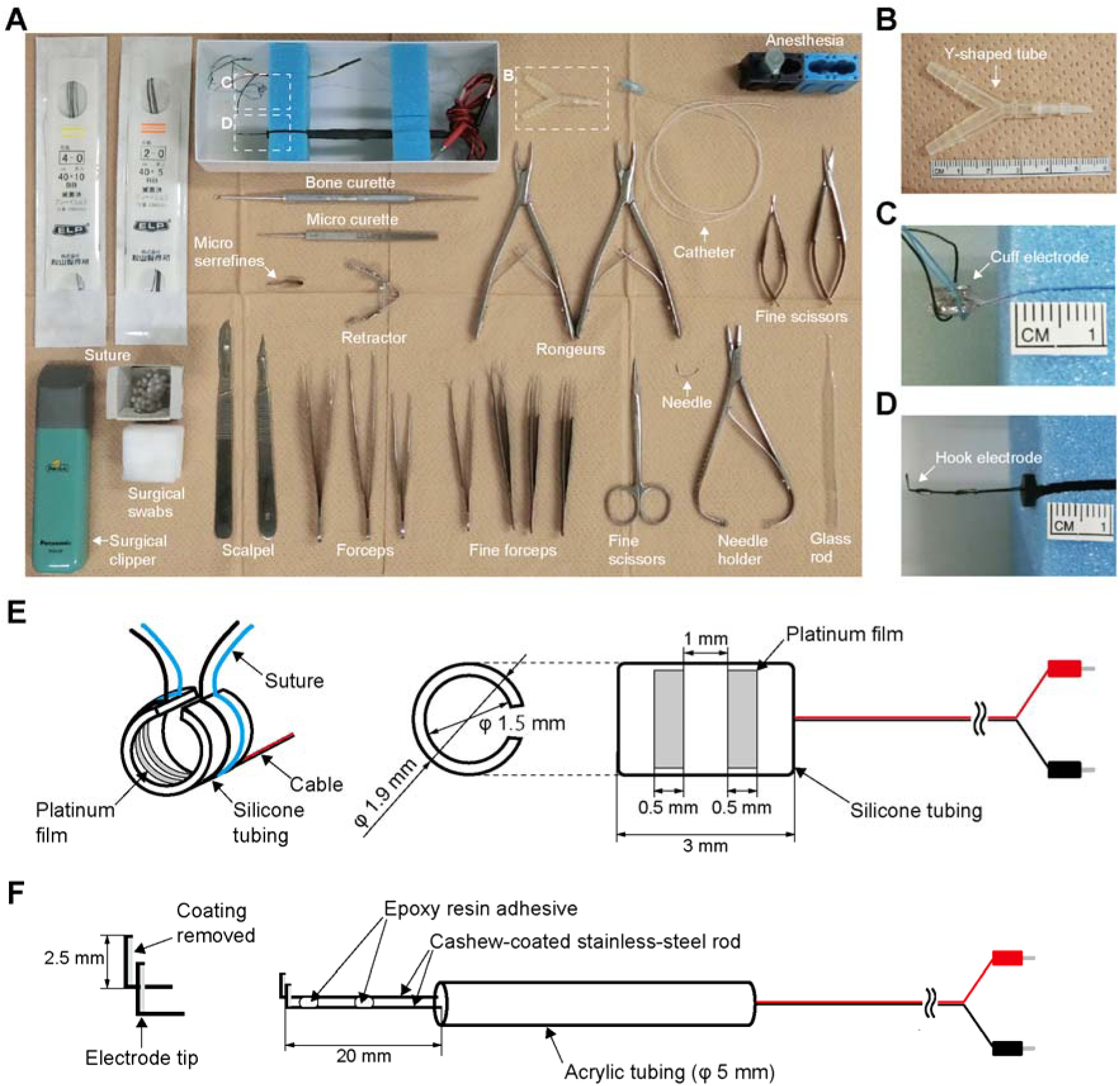
Acute experiment preparation. (A) Surgical tools, anesthesia, and consumables for the electrophysiological experiment. (B) Magnified view of the Y-shaped tube, as indicated by the square in (A). (C) Magnified view of the cuff electrode, as indicated by the square in (A). (D) Magnified view of the hook electrode, as indicated by the square in (A). (E) Specification diagram of the cuff electrode with dimensions. (F) Specification diagram of the hook electrode with dimensions.

## Key resources table

| REAGENT or RESOURCE | SOURCE | IDENTIFIER |
| --- | --- | --- |
| Bacterial and virus strains |  |  |
| AAV9-hSyn-ChR2(H134R)/EYFP | Kubota et al. 2019<sup>7</sup> | N/A |
| AAV9-CMV-eNpHR3.0/EYFP | Kosugi et al. 2025<sup>1</sup> | N/A |
| Chemicals, peptides, and recombinant proteins |  |  |
| Atipamezole |  | CAS#: 104054-27-5 |
| Butorphanol |  | CAS#: 42408-82-2 |
| Fast Green |  | CAS#: 2353-45-9 |
| Fluorinert |  | CAS#: 51142-49-5 |
| Isoflurane |  | CAS#: 26675-46-7 |
| Lidocaine |  | CAS#: 137-58-6 |
| Medetomidine |  | CAS#: 86347-14-0 |
| Midazolam |  | CAS#: 59467-70-8 |
| Mineral oil |  | CAS#: 8042-47-5 |
| Paraformaldehyde |  | CAS# 30525-89-4 |
| Phosphate-buffered saline |  | N/A |
| Rocuronium bromide |  | CAS#: 119302-91-9 |
| Saline |  | CAS#: 7647-14-5 |
| Thiopental sodium |  | CAS#: 71-73-8 |
| Experimental models: Organisms/strains |  |  |
| Rat: Wistar | Japan SLC, Inc. | N/A |
| Software and algorithms |  |  |
| Doric Neuroscience Studio | Doric Lenses | RRID: SCR_018569 |
| pClamp 11.2 | Molecular Devices | RRID: SCR_011323 |
| Other |  |  |
| Bone curette | YDM | 14-310 |
| Bone scissors | Hammacher | HSB648-10 |
| Fine forceps (curved) | FRIGZ | A049-0751 |
| Fine forceps (Dumont No. 5) | FST | 11254-20 |
| Fine forceps (Dumont No. 7) | FST | 11271-30 |
| Fine scissors | FST | 14060-11 |
| Fine scissors (Noyes Spring Scissors) | FST | 15011-12 |
| Fine scissors (Vannas Spring Scissors) | FST | 15000-00 |
| Forceps (Adson tissue forceps) | FRIGZ | A037-0524 |
| Forceps (Adson micro forceps) | FRIGZ | A045-0698 |
| Forceps (Semken tissue forceps) | FRIGZ | A037-0527 |
| Forceps (toothed Graefe forceps) | FST | 11053-10 |
| Hemostats (Mosquito hemostats) | FRIGZ | F052-0784 |
| Micro curette | FST | 10082-15 |
| Micro serrefines (Schwartz micro serrefines) | FST | 18052-01 |
| Needle holder (Mathieu needle holder) | FST | 12010-14 |
| Perfusion cannula | Natsume Seisakusho | KN-348-20G-80 |
| Retractor (Agricola retractor) | FST | 17005-04 |
| Rongeurs (Micro Friedman-Pearson Rongeurs) | FST | 16220-14 |
| Rongeurs (Micro Friedman-Pearson Rongeurs) | FST | 16221-14 |
| Scalpel handle (No.3) | FRIGZ | Z505-132 |
| Scalpel handle (No.4) | FRIGZ | Z505-136 |
| Surgical scissors | FRIGZ | B022-1306 |
| Absorbable suture (6-0) | Johnson & Johnson | D5890 |
| Braded silk suture (2-0) | Akiyama Seisakusho | N/A |
| Braded silk suture (4-0) | Akiyama Seisakusho | N/A |
| Catheter (3Fr) | Sansho | No.3 |
| Hamilton syringe (25 µL) | Hamilton | 1702RN |
| Heating pad |  | N/A |
| Microcentrifuge tube (1.5 mL) | Eppendorf | N/A |
| Micropipette (P10) | Gilson | F144055M |
| Myjector syringe (1 mL) |  | N/A |
| Non-absorbable suture (5-0) | Kono Seisakusho | NB085 |
| Paper towel |  | N/A |
| Pasteur pipette | Thermo Fisher Scientific | N/A |
| Polyethylene tubing with connector | Eicom | JT-10-80 |
| Scalpel blade (11) | FEATHER Safety Razor | N/A |
| Scalpel blade (21) | FEATHER Safety Razor | N/A |
| Self-closing forceps |  | N/A |
| Styrofoam |  | N/A |
| Surgical clipper |  | N/A |
| Surgical swab |  | N/A |
| Surgical tape |  | N/A |
| Syringe (1 mL) |  | N/A |
| Syringe (10 mL) |  | N/A |
| Syringe (50 mL) |  | N/A |
| Syringe needle (31G) |  | N/A |
| Timer |  | N/A |
| Three-way stopcock |  | N/A |
| Y-shaped tube |  | N/A |
| Amplifier | Nihon Kohden | MEG-6108M |
| Analog/digital converter | Molecular Devices | Digidata 1550 |
| Catherin needle (23G) |  | N/A |
| Cuff electrode | Unique Medical | Cat#TY-222-031d |
| Electrical stimulator | Nihon Kohden | SEN-7203 |
| Hook electrode | Unique Medical | Cat#TY-222-018a |
| Isoflurane vaporizer | Shinano Seisakusho | SN-487-OT Air+O<sub>2</sub> |
| Isolator | Nihon Kohden | SS-202J |
| Light source | Doric Lenses | LISER_LED470 |
| Manipulator | Narishige | SM-15L |
| Microinjection pump | Chemyx | Nanojet |
| Multimode optical fiber | Doric Lenses | MFP_960/1000/2200<br>-0.63_2m_FCM-ZF2.5 |
| Power mater | Thorlabs | PM100D |
| Pulse oximeter | Kent Scientific | MouseSTAT Jr. |
| Respirator | Shinano Seisakusho | SN-480-07 |
| Stereomicroscope | Konan Medical | POM 50- II |
| Stereotaxic frame | Narishige | SR-6R-HT |
| Thermometer | Physitemp | PTM1 |
| Weighing scale |  | N/A |

## Materials and equipment setup

1. Prepare the surgical tool used to isolate the nerve and the root: a glass rod with a bent tip. To make this tool. cut the tip of a Pasteur pipette and bend it to a 90-degree angle using a gas burner.
2. Prepare the stimulus electrode: a bipolar cuff electrode (Fig. 2C). It is recommended to use a silicone tube cuff with an inner diameter of 1.5 mm, equipped with platinum electrodes that are 0.5 mm wide and spaced 1 mm apart (Fig. 2E). We commissioned the fabrication to Unique Medical (Tokyo, Japan)
3. Prepare the recording electrode: a bipolar hook electrode (Fig. 2D). It is recommended to use a stainless-steel rod coated in cashew, with the tip bent at 2.5 mm and the coating removed to expose the electrode (Fig. 2F). We commissioned the fabrication to Unique Medical (Tokyo, Japan)

## Step-by-step method details

### Viral injection into the sciatic nerve

Timing: 1 h and 15 min

This position of the protocol details the procedure for intra-nerve AAV injection. It will be used to optogenetically modulate peripheral sensory nerve activity in later step.

1. Inject the rat intraperitoneally with 0.2 mg/kg medetomidine, 2.0 mg/kg midazolam, and 2.5 mg/kg butorphanol.
2. Verify adequate depth of anesthesia by monitoring pupil size and flexion reflex to the paw pinch.
3. Shave from the left gluteal region to the left knee.
4. Inject 0.2 mL of lidocaine subcutaneously into the left gluteal region for local anesthesia.
5. Put the animal on the heating pad and fixed in a prone position.
6. Begin monitoring heart rate, oxygen saturation, and rectal temperature.
7. Make a 10 mm incision approximately 5 mm to the left of the midline and 10 mm caudal to the iliac crest, directed toward the femoral head. (Fig. 3A).
8. Lift the gluteus maximus muscle and make a 10 mm incision parallel to the spine using fine scissors.
9. Open the incision site with a retractor to ensure adequate exposure (Fig. 3B).
10. Identify the sciatic nerve below the biceps femoris muscle. The sciatic nerve passes through the deeper layers of tissue.
11. Isolate the nerve approximately 20 mm from the surrounding tissue using the glass rod with a bent tip (Fig. 3C). ***Note*:** Please be careful, as a large blood vessel runs very close to the nerve.
12. Make a string from a surgical swab, wet it with saline, and pass it under the nerve.
13. Make a loop out of the string and clamp the ends with hemostat (Fig. 3D).
14. Apply tension to the nerve by lifting the string with hemostat. **CRITICAL:** Be careful not to apply excessive tension to the nerve, as this may cause damage. Also, ensure that the nerve does not become twisted, as this can increase the risk of leakage of the injected solution.
15. 15. Place a small surgical swab under the nerve to make it easier to check for leakage during injection (Fig. 3D).
16. 16. Insert a 31-gauge needle connected to a polyethylene tubing into the left side of the nerve (Fig. 3E).

**Figure 3.**
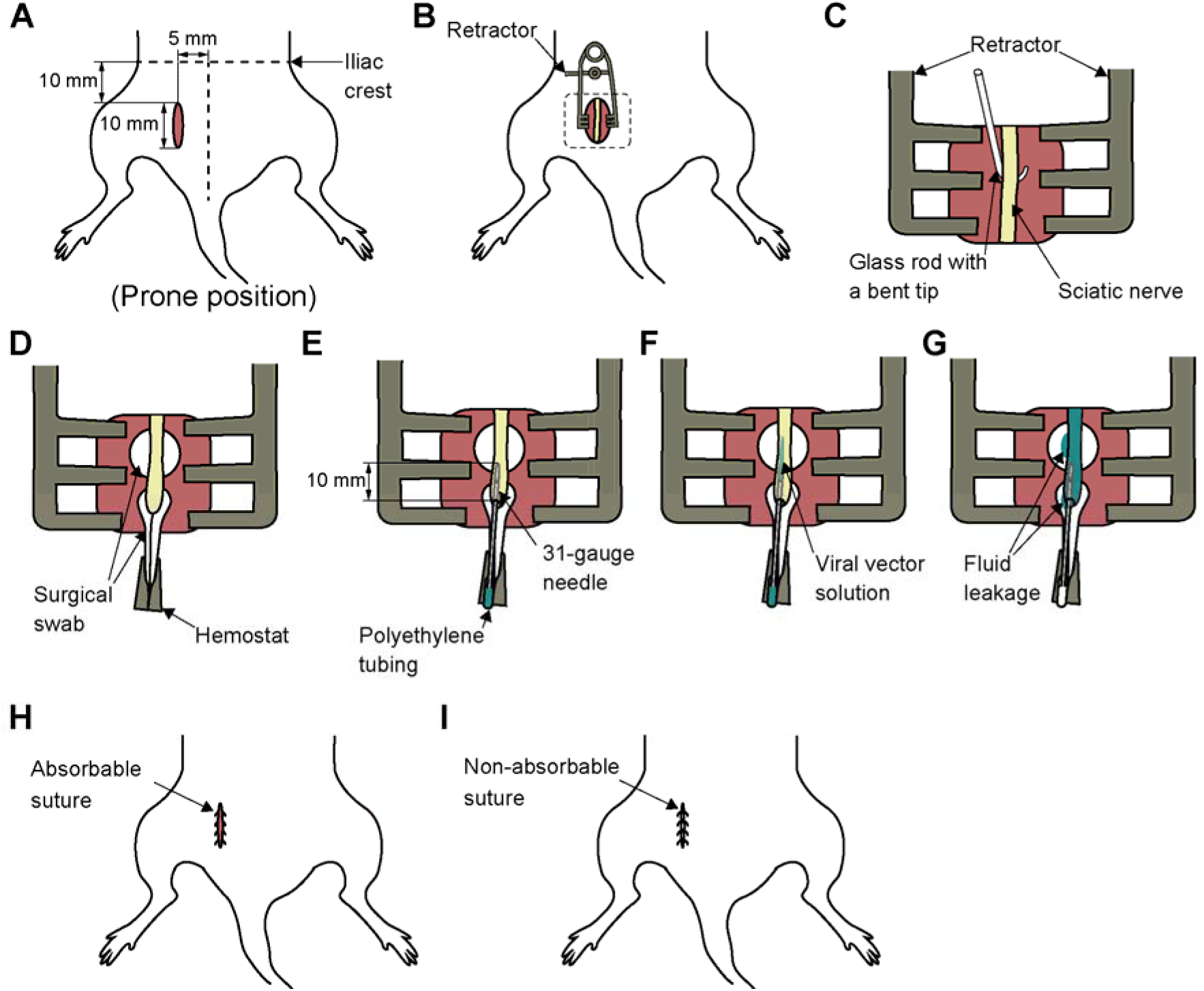
Detailed method for intra-nerve AAV injection. (A) Location of the incision. (B) Placement of a retractor. (C) Isolation of the sciatic nerve, as indicated by the square in (B). (D) Lifting the nerve with a surgical swab. (E) Insertion of a 31-gauge needle into the nerve. (F) Injection of the viral vector solution into the nerve. (G) Observation of the injection site. (H) Muscle closure with absorbable suture. (I) Skin closure with non-absorbable suture.

***Note*:** It is recommended to run a small amount of fluid flow before insertion to confirm that there is no clogging. The optimal insertion site is near the clamping area, preferably at the proximal end.

17. Advance the needle approximately 10 mm with the tip oriented upward, keeping it parallel to the nerve.

**CRITICAL:** If the needle is inserted too deeply, it may pass through the target area and cause leakage onto the swab placed underneath. If the insertion is too shallow, it may result in leakage from the insertion site. Re-insertion should be avoided as much as possible, as leakage tends to occur from the initial puncture hole.

18. After advancing the needle 10 mm, pull it back slightly to release the pressure.
19. Wait 5 minutes to allow tissue to seal around the needle tip.
20. Start the injection. Inject 3 μL of viral vector solution at a rate of 0.6 μL/min (Fig. 3F).

***Note*:** During the injection, the needle should not be fixed but allowed to float. In addition, be sure to check for any fluid leakage onto the swab placed underneath.

21. Wait 10 minutes after the end of the injection to ensure absorption of the solution.

***Note*:** To prevent leakage of the injected solution after the surgery, be sure to wait for 10 minutes.

22. Remove the needle.
23. Insert a 31-gauge needle into the right side of the nerve.
24. Repeat steps 17-22

***Note*:** Two separate injections were performed into the common peroneal and tibial branches of the sciatic nerve to ensure that the nerve was filled uniformly (Fig. 3G). In total, 6 μL of viral vector solution was injected.

25. Irrigate around the injection site.
26. Close the muscle with an absorbable suture (Fig. 3H).
27. Close the skin with a non-absorbable suture (Fig. 3I).
28. Inject the antagonist of anesthesia and allow to recover at 37°C.

### Preparation for in vivo dorsal root recording

Timing: 5 h

Timing: 4 weeks after steps 1-28

This position of the protocol details the procedure for preparation of dorsal root potential recording at the lumber level.

29. Inject the rat intraperitoneally with 0.2 mg/kg medetomidine, 2.0 mg/kg midazolam, and 2.5 mg/kg butorphanol.
30. Verify adequate depth of anesthesia by monitoring pupil size and flexion reflex to the paw pinch.
31. Shave the back (from the caudal end of the sternum to the tail), the ventral surface (from the larynx to the abdomen), and both thighs (ventral and dorsal sides).
32. Put the animal on the heating pad and fixed in a supine position.

***Note*:** Position the head as flat as possible to keep the trachea horizontal. If necessary, place a paper towel or similar support under the neck to maintain alignment.

33. Bind both hands and feet.
34. Begin monitoring heart rate, oxygen saturation, and rectal temperature.
35. Start tracheotomy. First, make a 20 mm incision in the skin above the trachea, at the level between the ear and the shoulders (Fig. 4A).
36. Expose and isolate the trachea from the surrounding tissue (Fig. 4B).
37. Lift the trachea and pass two 2-0 braided silk sutures underneath it (Fig. 4C).
38. Make an incision in the trachea between the two sutures (Fig. 4 D).

**Figure 4.**
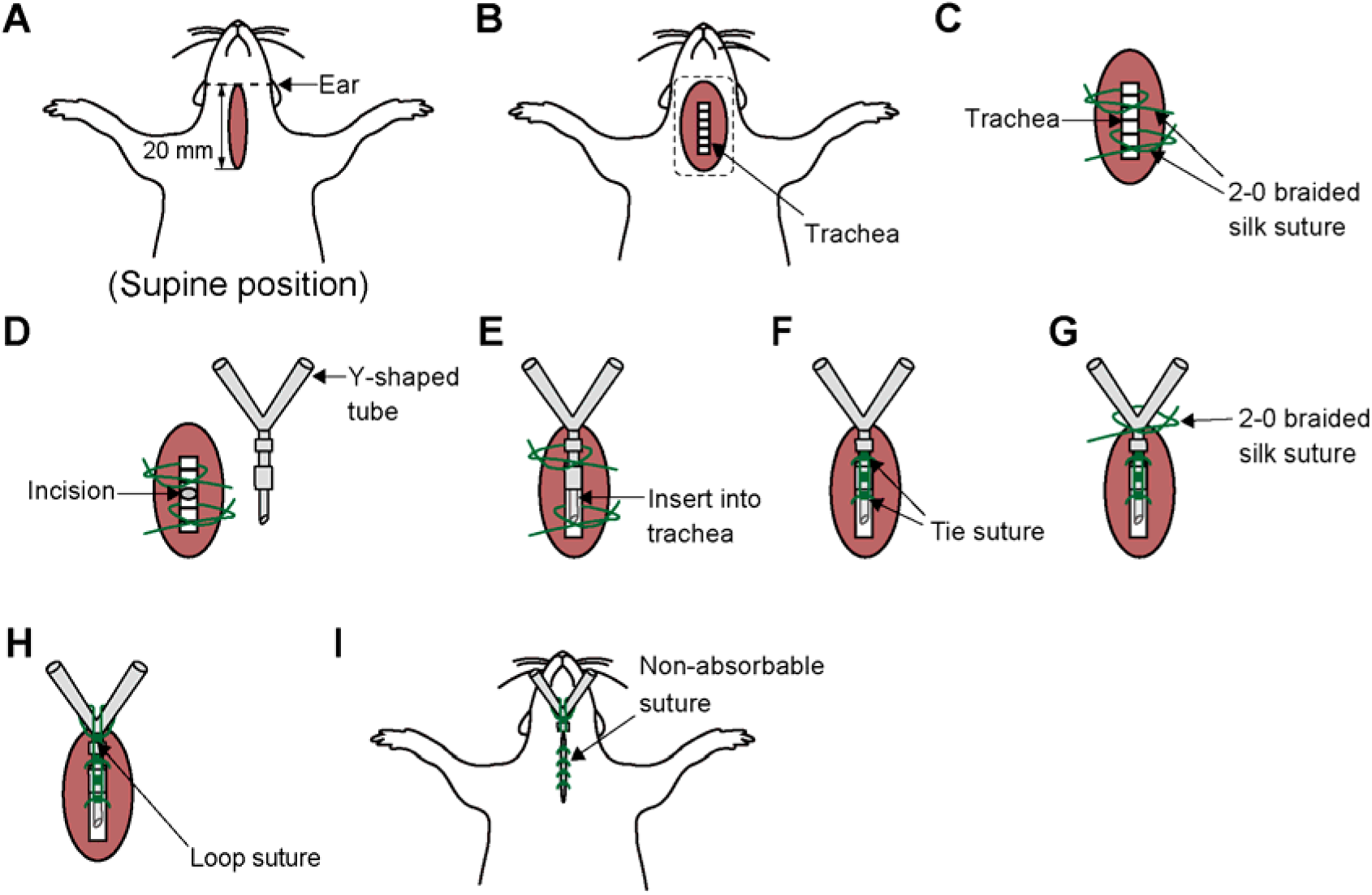
Detailed method for tracheotomy. (A) Location of the incision. (B) Exposure of the trachea (C) Isolation of the trachea, as indicated by the square in (B). (D) Making an incision in the trachea. (E) Insertion of the Y-shaped tube into the trachea. (F) Securing the tube by tying the suture. (G) Further securing the tube by tying the suture. (H) Placement of a loop suture for stabilization. (I) Skin closure with non-absorbable suture.

***Note*:** Take care to prevent blood or other fluids from entering the trachea through the incision site. It is recommended to gently lift the trachea by holding up 2-0 braided silk sutures placed underneath it.

39. Insert a Y-shaped tube into the tracheal incision (Fig. 4 E).

***Note*:** To prevent the tip from contacting the tracheal wall and causing an obstruction, ensure that the Y-tube is inserted horizontally.

40. Tie the sutures around the trachea to secure the Y-shaped tube (Fig. 4F).

***Note*:** Tie the caudal suture first, then tie the rostral suture. Finally, tie the two sutures together.

41. Place and tie a 2-0 braided silk suture to the skin at the bifurcation of the Y-shaped tube, then wrap it around the tube to further secure it (Fig. 4G).
42. Loop the suture around the bifurcation again and tie it to the previously placed sutures (Fig. 4H).
43. Close the skin with a non-absorbable suture (Fig. 4I).
44. Connect the Y-shaped tube to the ventilator.
45. Start mechanical ventilation.

***Note*:** Carefully monitor for synchronization between chest movements and ventilator activity.

46. Switch the anesthesia to 0.5–1% isoflurane.
47. Begin establishing an intravenous line. First, make a 20 mm incision on the medial thigh skin (Fig. 5A).
48. Open the incision site with a retractor to ensure adequate exposure (Fig. 5B).
49. Expose and isolate the femoral vein from the surrounding tissue (Fig. 5C).

**Figure 5.**
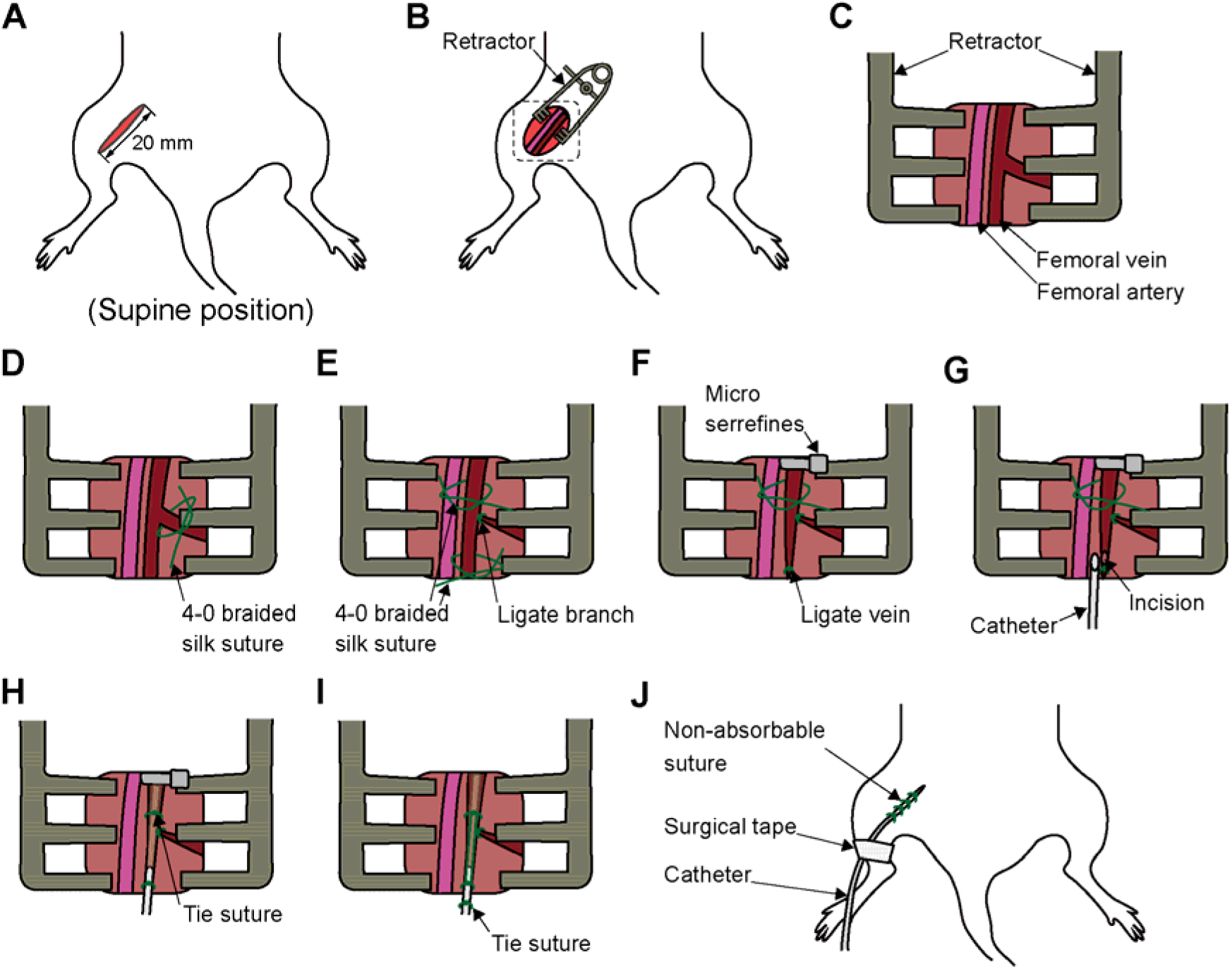
Detailed method for establishing an intravenous line. (A) Location of the incision. (B) Placement of a retractor. (C) Exposure of the femoral vein, as indicated by the square in (B). (D) Isolation of the vein. (E) Ligation of the vein branch. (F) Ligation of the vein using micro serrefines. (G) Insertion of the catheter. (H) Securing the catheter by tying the suture. (I) Further securing the catheter by tying the suture. (J) Skin closure with non-absorbable suture.

***Note*:** When exposing the femoral vein, dissect as proximally as possible. The vein is thicker closer to the proximal end, making catheter insertion easier. Take care not to damage the artery, as it runs immediately adjacent to the vein.

50. Ligate the femoral vein branches (Fig. 5D).

***Note*:** The more branches are ligated, and a straight segment is ensured, the easier it becomes to insert the catheter.

51. Lift the femoral vein and pass two 4-0 braided silk sutures underneath it (Fig. 5E).

***Note*:** Take care to avoid twisting the vein when passing the 4-0 braided silk suture under the vein.

52. Clamp the femoral vein proximal to the sutures using a micro serrefine (Fig. 5F).

***Note*:** Clamp the proximal end first to engorge the vessel.

53. Ligate the distal portion of the femoral vein with a suture (Fig. 5F).
54. Make an incision just proximal to the ligature and insert a catheter (Fig. 5G).

***Note*:** Grasp the vessel incision with fine forceps and use curved fine forceps to handle the catheter. If the catheter cannot be inserted smoothly, withdraw it slightly. Failure to do so may result in vessel rupture. If the clamp is loose or a branch is overlooked, blood may continue to flow. In such cases, ligate the suture on the proximal side immediately to stop the blood flow.

55. Tie the sutures around the vein to secure the catheter (Fig. 5H).

***Note*:** Tie the proximal suture first, then tie the distal suture. Finally, tie the two sutures together.

56. Place and tie a 2-0 braided silk suture to the skin at the distal end of the incision, then wrap it around the catheter to further secure it (Fig. 5I).
57. Slightly withdraw the syringe to check for blood backflow.
58. Close the skin with a non-absorbable suture (Fig. 5J).
59. Secure the catheter to the thigh with surgical tape to prevent dislodgement (Fig. 5J).
60. Turn the animal over and place it in the prone position.

***Note*:** It is recommended to temporarily remove the electrocardiogram, oxygen saturation, and rectal temperature probes as well as the ventilator tubing.

61. Begin the laminectomy. First make an incision in the skin from just above the floating rib to just above the iliac crest (Fig. 6A).
62. Expose the spinous processes from the 13th thoracic vertebra (Th13) to the 1st sacral vertebra (S1) by dissecting the erector spinae muscles (Fig. 6B).

**Figure 6.**
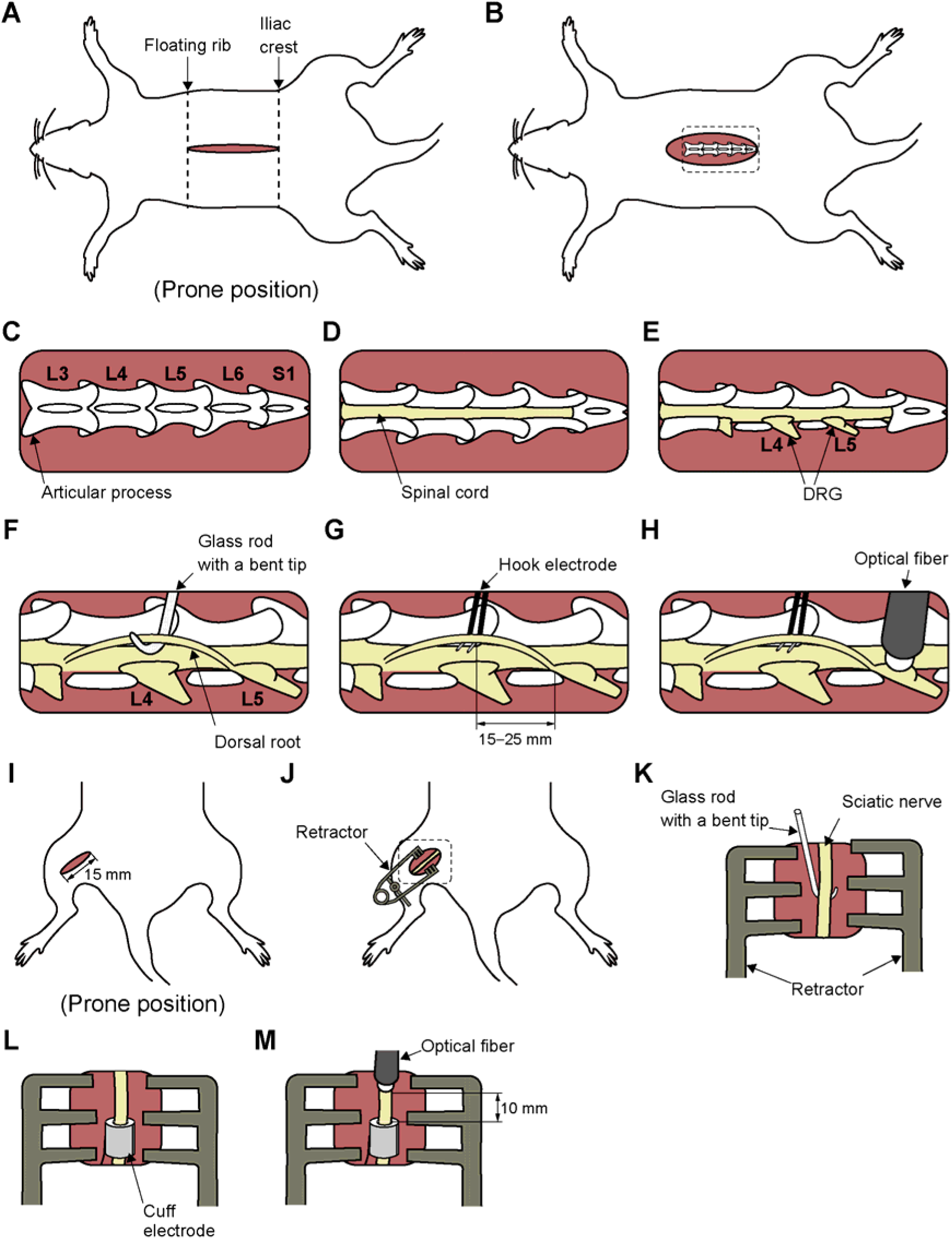
Detailed method for laminectomy. (A) Location of the incision. (B) Exposure of the spinous processes. (C) Exposure of the articular processes, as indicated by the square in (B). (D) Exposure of the spinal cord by laminectomy. (E) Exposure of the dorsal root ganglia (DRG). (F) Isolation of the dorsal root. (G) Mounting a bipolar hook electrode. (H) Placement of a multimode optical fiber above the DRG. (I) Location of the incision. (J) Placement of a retractor. (K) Isolation of the sciatic nerve, as indicated by the square in (J). (L) Mounting a bipolar cuff electrode. (M) Placement of the optical fiber above the nerve.

***Note*:** Segmental levels are determined based on vertebral mobility. Vertebrae below S1 lack intervertebral motion and are thus considered immobile.

63. Expose the articular processes using fine scissors, bone curette, and micro curette (Fig. 6C).
64. Perform a laminectomy from the 1st lumbar vertebra (L1) to S1 (Fig. 6D).

***Note*:** Holding the vertebral body with a toothed forceps and creating sufficient space while maintaining spinal extension allows safe insertion of a bone rongeur without risking damage to the spinal cord.

65. Expose the 4th (L4) and 5th (L5) lumbar DRGs (Fig. 6E).

**CRITICAL:** The region around the DRG is particularly prone to bleeding, which can obstruct light delivery and prevent the recording of responses to DRG irradiation. Sufficient lateral exposure is recommended to minimize blood pooling over the DRG.

66. Begin nerve exposure. Make a 15 mm incision on the lateral thigh skin and identify the sciatic nerve between the vastus lateralis and the biceps femoris muscles (Fig. 6I).
67. Expose and isolate the sciatic nerve approximately 20 mm from the surrounding tissue using the glass rod with a bent tip (Fig. 6J, K).
68. Fix the animal in the stereotaxic frame.

***Note*:** Make sure the endotracheal tube and the intravenous catheter do not become dislodged while securing the animal in the stereotaxic frame. After securing the fixation, check the ventilation status again. It is recommended to temporarily remove the electrocardiogram, oxygen saturation, and rectal temperature probes.

69. Suspend the limbs on the stereotaxic frame to keep the body elevated.
70. Clamp both the upper thoracic spine and the base of the tail to immobilize the animal.
71. Raise the skin and muscle around the exposed DRG and nerve and tie them to form a pool filled with mineral oil to protect the exposed DRG and nerve from drying out.

***Note*:** Warm the mineral oil to 36–40 °C before use, and periodically check the temperature and volume, as both may decrease over time.

72. Switch the anesthesia to intravenous administration of thiopental sodium (10 mg/kg).
73. Administer rocuronium bromide intravenously (10 mg/kg) to achieve neuromuscular blockade.
74. Open the dura with a sharp needle.
75. Identify the L4 and L5 dorsal roots and separate them from the surrounding root using the glass rod with a bent tip (Fig. 6F).

***Note*:** To prevent drying out, perform this procedure in the mineral oil.

### Recording dorsal root potentials

Timing: 4 h

This position of the protocol details the procedure for recording dorsal root potentials evoked by electrical or optical stimulation of the sciatic nerve.

76. Mount a bipolar cuff electrode on the isolated sciatic nerve to apply electrical stimulation (Fig. 6L).
77. Place a multimode optical fiber 10 mm proximal to the cuff electrode (Fig. 6M).

***Note:*** The tip of the optical probe was positioned perpendicular to the nerve so that the diameter of the irradiation area was approximately equal to the diameter of the probe. At this stage, if a blood clot covers the nerve, remove it before placing the optical fiber. Light transmission may be obstructed by bleeding, resulting in reduced effectiveness of optical stimulation.

78. Mount a bipolar hook electrode on the isolated L4 or L5 dorsal root at 15–25 mm proximal to DRG to record stimulus-evoked volleys (Fig. 6G).

***Note*:** Avoid lifting the dorsal root excessively.

79. Put a ground electrode on the exposed back muscle.
80. Start recording.

***Note*:** If optical stimulation of the DRG is desired, place the optical fiber just above DRG (Fig. 6H). In this case, the tip of the optical probe was positioned perpendicular to the surface of DRG so that the diameter of the irradiation area was approximately equal to the diameter of the probe. At this stage, if a blood clot covers the DRG, remove it before placing the optical fiber. Light transmission may be obstructed by bleeding, resulting in reduced effectiveness of optical stimulation.

81. At the end of the experiments, annotate stimulus and recording sites. Make small electrolytic lesions in the dorsal root and sciatic nerve using a 40 μA direct current through the recording and stimulating electrodes for 40 s.
82. Start perfusion. First, deeply anesthetize the animal via intravenous administration of thiopental sodium (20 mg/kg).
83. Perfuse transcardially with 50 mL of phosphate-buffered saline (PBS; pH 7.4), followed by 300 mL of 4% paraformaldehyde using a perfusion cannula.
84. After perfusion, remove the lumbar region of the spinal cord together with the DRG and sciatic nerve.

## Expected outcomes

Begin by confirming the response to electrical stimulation. When electrical stimulation is applied to the sciatic nerve via a bipolar cuff electrode, the evoked incoming afferent volleys can be recorded at the dorsal root (Fig. 7A). The size of the evoked volley should vary with stimulus intensity, with its amplitude increasing rapidly at lower currents and more gradually at higher currents (Fig. 7B).

**Figure 7.**
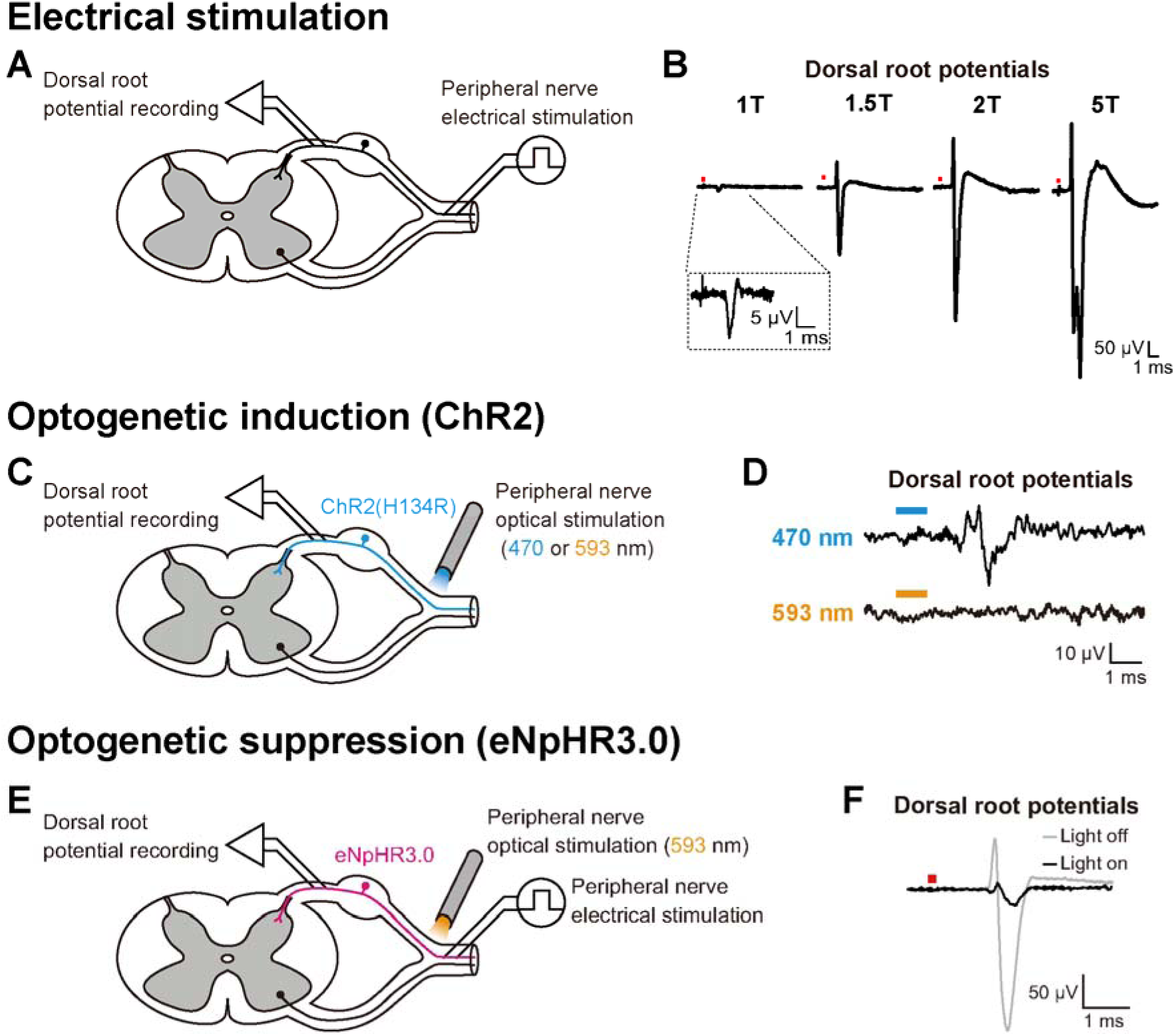
Electrophysiological assessment of optogenetic modulations of sensory afferent activity. (A) Schema of the acute *in vivo* electrophysiological experiment. (B) Examples of electrically evoked dorsal root potentials using different intensities of electrical stimulation. Waveforms represent the stimulus-triggered averaging of 20 responses. The red square above the waveforms indicates the timing of electrical stimulation. (C) Schema of the acute *in vivo* electrophysiological experiment following channelrhodopsin (ChR2) transduction. (D) Examples of dorsal root potentials evoked by light stimulation with different wavelengths. Waveforms represent the stimulus-triggered averaging of 20 responses. The bar above each waveform indicates the timing of light irradiation. (E) Schema of the acute *in vivo* electrophysiological experiment following halorhodopsin (eNpHR3.0) transduction. (F) Examples of electrically evoked dorsal root potentials with (black line) and without (gray line) light irradiation. Waveforms represent the stimulus-triggered averaging of 20 responses. The red square above the waveforms indicates the timing of electrical stimulation.

Electrical nerve stimulation recruits action potentials depending on axonal diameter. Axons with larger diameters exhibit lower electrical resistance and are therefore more easily depolarized by weaker input currents^8,9^. The earliest component of the recorded volleys, termed the “Aα” wave, represents action potentials generated by group I muscle afferent fibers. These fibers possess the largest axonal diameters among the different classes of primary afferents.

In rats transduced with ChR2, when blue light is delivered to the sciatic nerve through the optical fiber (Fig. 7C), evoked volleys can be recorded at the dorsal root (Fig. 7D, top row), whereas yellow light should not produce any detectable effect (Fig. 7D, bottom row). The maximum amplitude of the optically evoked volleys, normalized to the amplitude of the electrically evoked volleys, is typically around 4.3% (range, 1.4–20%; *n* = 7). This value represents the expected strength of optogenetic activation under the conditions described in this protocol.

In rats transduced with eNpHR3.0, when the sciatic nerve was pre-activated by electrical stimulation through a bipolar cuff electrode mounted onto the nerve at a fixed stimulus intensity (1.5 times as high as the threshold intensity), and yellow light was applied proximally (Fig. 7E), the volley size evoked by electrical stimulation (Fig. 7F, gray line) was diminished (Fig. 7F, black line). The maximum percentage of optogenetic inhibition, calculated by normalizing the amplitude of the electrically evoked volleys under the light-on condition to that under the light-off condition, is typically around 26% (range, 13–85%; *n* = 10). This value represents the expected strength of optogenetic inhibition under the conditions described in this protocol.

## Limitations

The injection parameters, such as the injection volume and speed, and the titer of the viral vectors, were optimized for the sciatic nerve of 4-week-old male rats. When applying this protocol to different nerves, sexes, or species, future work should be performed to optimize parameters.

Furthermore, it should be noted that when extrapolating this method to other species, target specificity may differ even when using the same AAV serotype and injection approach. We previously reported that intra-nerve injection of AAV9 resulted in different target specificity to DRG neurons between rats and marmosets, with selective transduction of large-diameter DRG neurons observed in rats but not clearly in marmosets^2^. Therefore, species-specific validation of AAV transduction properties is necessary when adapting the protocol to other animal models.

## Troubleshooting

### Problem 1

The injected viral solution leaks onto the cotton swab beneath the nerve.

### Potential solution

Check the site of leakage (Step 20). If leakage is observed near the needle insertion site, it may indicate that the needle is inserted too shallowly; in this case, advance the needle slightly. If leakage appears away from the insertion site, the needle may be inserted too deeply or may be exerting tension on the nerve bundle. In such cases, slightly withdraw the needle or adjust its position so that the needle tip is aligned parallel to the nerve bundle.

### Problem 2

The injected viral solution becomes clogged and does not come out of the tip of the needle.

### Potential solution

Be sure to check for any clogging before inserting the needle (Step 16). In some cases, even if there is no clogging before insertion, the needle may become clogged within the nerve bundle. In such situations, pulling the needle back slightly may result in a sudden outflow of fluid (Step 18). If the clogging still does not resolve, stop the syringe pump temporarily.

### Problem 3

Unstable vital monitor.

### Potential solution

- Before beginning the experiment, confirm that there are no obstructions or foreign materials in the tracheal tube or intravenous catheter (Steps 9 and 10). At the same time, ensure that the ventilation rate and tidal volume are appropriately set.
- Throughout the experiment, continuously monitor for possible tracheal tube obstruction. If the tube becomes blocked and internal pressure increases, ventilation may be compromised. An increase in pressure is often identified by excessive expansion of the back during ventilation.
- If ventilation problems occur after stereotaxic fixation, such as desynchronization between chest movements and ventilator activity, immediately remove the animal from the stereotaxic frame (Step 68). Then, disconnect the Y-shaped tube from the ventilator, block one branch of the tube, and forcefully deliver air through the other branch using a blower or equivalent.
- If bradycardia (a decrease in heart rate) occurs for reasons unrelated to the ongoing procedure, temporarily reducing the ventilation rate may help. Confirm whether this adjustment leads to a recovery in heart rate.

### Problem 4

Difficulty in recording dorsal root potentials.

### Potential solution

- If clear electrically-evoked dorsal root potentials are not observed (see Fig. 8A for a favorable example), there may be issues with either the stimulation or the recording processes. First, to check for stimulation problems, apply supramaximal electrical stimulation (approximately > 1 mA).
- If the issue lies with the recording, there are two possible causes: either excessive noise resulting in a low signal-to-noise ratio, or small response amplitudes. If the noise level is excessive (baseline fluctuations exceeding ±50 µV), ensure that the electrodes are properly immersed in the oil pool (Step 71), and check this regularly, as the oil level may decrease over time. It may also help to reposition the ground electrode (Step 79) or to place the amplifier and isolator farther away from the animal.
- If both the noise level and the response amplitude are small, examine the waveform shape first. If a polyphasic waveform is observed (see Fig. 8A for an unfavorable example), it is likely that the nerve or root is damaged. Take care not to damage or excessively stretch the nerve during injection (Step 11) or preparation (Step 67). Similarly, avoid damaging the root during exposure (Steps 64 and 75), and do not lift it excessively with the hook electrode (Step 78).
- If the waveform is not polyphasic, the recorded dorsal root segment may be inaccurate (Step 75). Note that although there is no significant difference in amplitude between L4 and L5, the evoked response amplitude to sciatic nerve stimulation is smaller at L6.

**Figure 8.**
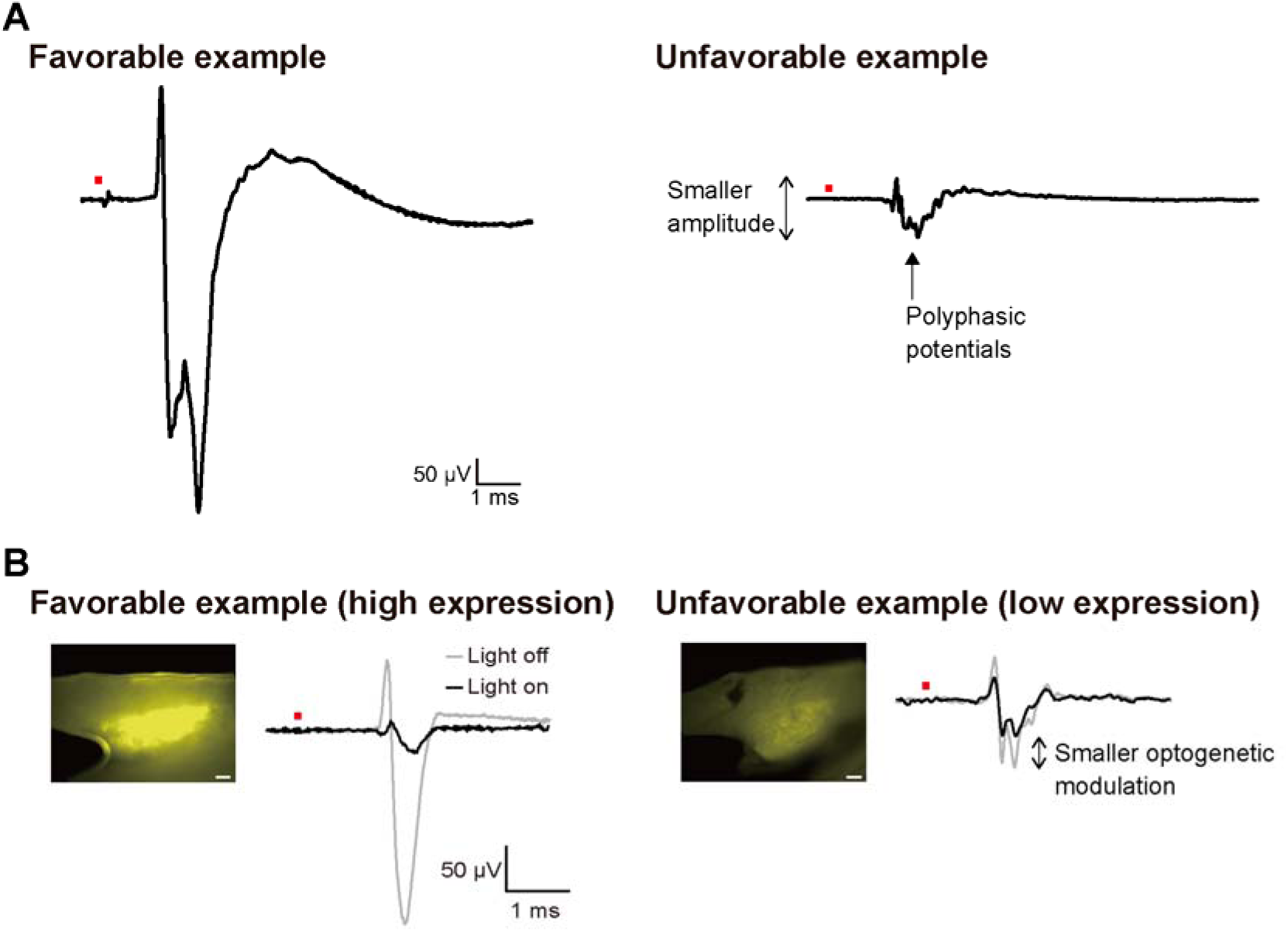
Examples of dorsal root potentials illustrating the range of possible outcomes, from favorable to unfavorable. (A) Representative examples of dorsal root potentials evoked by supramaximal electrical stimulation. The favorable example shows a large-amplitude potential, whereas the unfavorable example shows smaller amplitude and polyphasic potentials. Waveforms represent the stimulus-triggered averaging of 20 responses. The red square above the waveforms indicates the timing of electrical stimulation. (B) Relationship between opsin expression level and optogenetic modulation. Macroscopic images show the expression of native yellow fluorescent protein (YFP) in the entire DRG with the same detector value. In the favorable example (high expression), strong YFP fluorescence in the DRG (left) corresponds to large optogenetic modulation (right, black line: light on; gray line: light off). In the unfavorable example (low expression), weaker YFP fluorescence corresponds to smaller optogenetic modulation. Waveforms represent the stimulus-triggered averaging of 20 responses. The red square above the waveforms indicates the timing of electrical stimulation. Scale bars: 200 µm.

### Problem 5

Difficulty in confirming optogenetic effect.

### Potential solution

- If no effect is observed with optical stimulation despite a successful electrical stimulation response, the issue may lie in the level of opsin expression or the method of light delivery.
- First, check the opsin expression level. This can be assessed by examining the fluorescent signal intensity of YFP in the macroscopic image of the entire DRG before sectioning (Fig. 8B). Both excessively high and insufficient levels of opsin expression can impair the optogenetic effect. Excessive expression may lead to neuronal toxicity. Specifically, ChR2 has been reported to induce an immune response in the peripheral nervous system, resulting in loss of expression over time^10^. In such cases, the electrically evoked volleys also exhibit low amplitudes, as shown in Fig. 8A for an unfavorable example. Conversely, insufficient expression diminishes optogenetic effect, as the extent of optogenetic modulation correlates with the gene expression level (see Fig. 8B for an unfavorable example).
- If the expression level is sufficient, the issue may lie in light delivery. First, check whether light transmission is obstructed by bleeding or other factors around the irradiation site (Step 77). Next, confirm that the irradiation site is appropriate (Step 77). During this process, determine the optimal position while monitoring the response.

## Resource availability

### Lead Contact

Further information and requests for resources and reagents should be directed to, and will be fulfilled by, the lead contact, Kazuhiko Seki.

### Technical Contact

Questions about the technical specifics of the protocol should be directed to and will be fulfilled by the technical contact, Akito Kosugi

### Materials availability

The materials used in this study are available upon reasonable request.

### Data and code availability

This study did not generate or analyze data or code.

## Acknowledgments

We thank Dr. Ken-ichi Inoue for providing the AAVs used in this study, and Ms. Moeko Kudo for her technical assistance. This work was supported by a Grant-in-Aid from the Japan Society for the Promotion of Science (JSPS) (grant numbers JP19H05724, JP19H01092, JP26120003, and JP23H05488 [to K.S.], and JP21K17633 [to A.K.]), and research grants from the Japan Agency for Medical Research and Development (AMED) (grant numbers JP19ek0109216, JP21dm0207092, JP21dm0207066, JP24gm0010009, and JP21dm0207077 [to K.S.]). This work was also supported by the Collaborative Research in Computational Neuroscience (CRCNS) US–Japan Program of the National Science Foundation (NSF) (grant number 2113096 [to K.S.]).

## Author contributions

A.K. and K.S. wrote the manuscript. A.K., W.S., and S.K. generated the protocols included in the paper.

## Declaration of generative AI and AI-assisted technologies in the manuscript preparation process

During the preparation of this work the authors used ChatGPT (OpenAI) in order to assist with language editing and grammar correction. After using this tool, the authors reviewed and edited the content as needed and take full responsibility for the content of the published article.

## References

1. Kosugi, A., Kudo, M., Inoue, K., Takada, M., and Seki, K. (2025). Bidirectional optogenetic modulation of peripheral sensory nerve activity: Induction vs. suppression through channelrhodopsin and halorhodopsin. iScience. 28, 112178.

2. Kudo, M., Sidikejiang, W., Fujiwara, M., Saito, Y., Kubota, S., Inoue, K., Takada, M., and Seki, K. (2021). Specific gene expression in unmyelinated dorsal root ganglion neurons in nonhuman primates by intra-nerve injection of adeno-associated virus 6 vector. Mol. Ther. Methods Clin. Dev. 23, 11–22.

3. Deisseroth, K. (2011). Optogenetics. Nat. Methods 8, 26–29.

4. Deisseroth, K. (2015). Optogenetics: 10 years of microbial opsins in neuroscience. Nat. Neurosci. 18, 1213–1225.

5. Fischer, G., Kostic, S., Nakai, H., Park, F., Sapunar, D., Yu, H., and Hogan, Q. (2011). Direct injection into the dorsal root ganglion: Technical, behavioral, and histological observations. J. Neurosci. Methods 199, 43–55.

6. Yang, O.J., Robilotto, G.L., Alom, F., Alemán, K., Devulapally, K., Morris, A., and Mickle A.D. (2023). Evaluating the transduction efficiency of systemically delivered AAV vectors in the rat nervous system. Front. Neurosci. 17, 1001007.

7. Kubota, S., Sidikejiang, W., Kudo, M., Inoue, K., Umeda, T., Takada, M., and Seki, K. (2019). Optogenetic recruitment of spinal reflex pathways from large-diameter primary afferents in non-transgenic rats transduced with AAV9/Channelrhodopsin 2. J. Physiol. 597, 5025–5040.

8. McNeal, D.R. (1976). Analysis of a model for excitation of myelinated nerve. IEEE Trans. Biomed. Eng. 23, 329–337.

9. Lertmanorat, Z., and Durand, D.M. (2004). Extracellular voltage profile for reversing the recruitment order of peripheral nerve stimulation: A simulation study. J Neural. Eng. 1, 202–211.

10. Maimon, B.E., Diaz, M., Revol, E.C.M., Schneider, A.M., Leaker, B., Varela, C.E., Srinivasan, S., Weber, M.B., and Herr, H.M. (2018). Optogenetic Peripheral Nerve Immunogenicity. Sci. Rep. 8, 14076.

